# An Evolving-Dynamic Network Activity Approach to Epileptic Seizure Prediction using Machine Learning

**DOI:** 10.1101/2020.11.29.402461

**Authors:** Christine Joy Liu, Jordan Sorokin, Surya Ganguli, John Huguenard

## Abstract

Absence epilepsy is a neurological condition characterized by abnormally synchronous electrical activity within two mutually connected brain regions, the thalamus and cortex, that results in seizures and affects more than 6.5 million people. Epilepsy is commonly studied through the use of the electroencephalogram (EEG), a device that monitors brain waves over time. In this study, we introduced machine learning models to predict epileptic seizures in two ways, one to train logistic regression models to provide an accurate decision boundary to predict based off frequency features, and second to train convolutional neural networks to predict based off spectral power images from EEG. This pipeline employed a two model approach, using logistic regression and convolutional neural networks to predict seizures. The evaluation, performed on data from 9 mice, achieved prediction accuracies of 98%. The proposed methodology introduces a novel aspect of looking at predicting absence seizures, which are known to be short events, in addition to the comparison between a time-dependent and time-agnostic seizure prediction classifier. The overall goal of these experiments were to build a model that can accurately predict whether or not a seizure will occur.

## 1 Introduction and Objectives

Epilepsy is the 4th most common neurological condition affecting more than 65 million people. Absence epilepsy, a particular form of epilepsy, is a neurological disorder affecting children between the ages of 4 and 12, and accounts for approximately 10% of all patients with epilepsy. Although absence seizures have traditionally been described as stochastic events [1], recent research has discovered that the cortex and thalamus show notable changes in neural activity prior to seizure onset [2], which may predict seizure severity [3]. These pre-seizure changes demonstrate a dynamic shift in brain state associated with an evolution into a seizure, and are clinically useful as patients may be warned of an impending seizure prior to its initiation [4]. Currently, there is no model that can accurately predict epileptic seizures with sufficient warning time to administer anti-convulsant medications or relocate the patient to a safe location [5]. While this work demonstrates a link between pre-seizure brain state and the subsequent seizure, the relationship between the likelihood of seizures at any moment and the state of the brain remains unknown in general. Indeed, seizures tend to cluster in groups separated by long periods of normal brain activity as shown by preliminary work in the Huguenard Lab [3]. This suggest that, in addition to the pre-seizure period of each seizure, there is a slower shift in network state (brain state) that influences seizure probability [6]. It is crucial to understand this process in more detail as the potential power of predicting future seizure likelihood and severity is clinically invaluable and may vastly improve the well-being of epilepsy patients [7].

Current literature has focused on a binary classification approach between pre-seizure and seizure signals of an electroencephalogram (EEG), which is an electrophysiological monitoring method consisting of electrodes used to record electrical activity of the brain [8]. The standard approach is gathering data from EEG segments and splitting the data into two categories (seizure and non-seizure). We can then train machine learning classifiers to predict seizures by distinguishing between these pre-seizure, termed preictal, and non-seizure, termed interictal, segments. One of the biggest problem that patients face with spontaneously occurring seizures is being unsure when their seizures will occur [5]. Thus, having a better understanding of how early we can tell these predictions could be incredibly helpful for both patients and physicians.

In this paper, we introduce a new absence seizure prediction model built on the finding that there exists a longer and slower shift in network activity that influences probability of seizures. We implemented both a logistic regression classifier and convolutional neural networks to achieve high performance with predicting seizures. In addition, we explore the extent of the prediction window by varying the amount of time before seizure onset. By applying these existing machine learning models to a new form of epilepsy, we can attempt to better understand the mechanisms of absence seizures.

**Figure 1:**
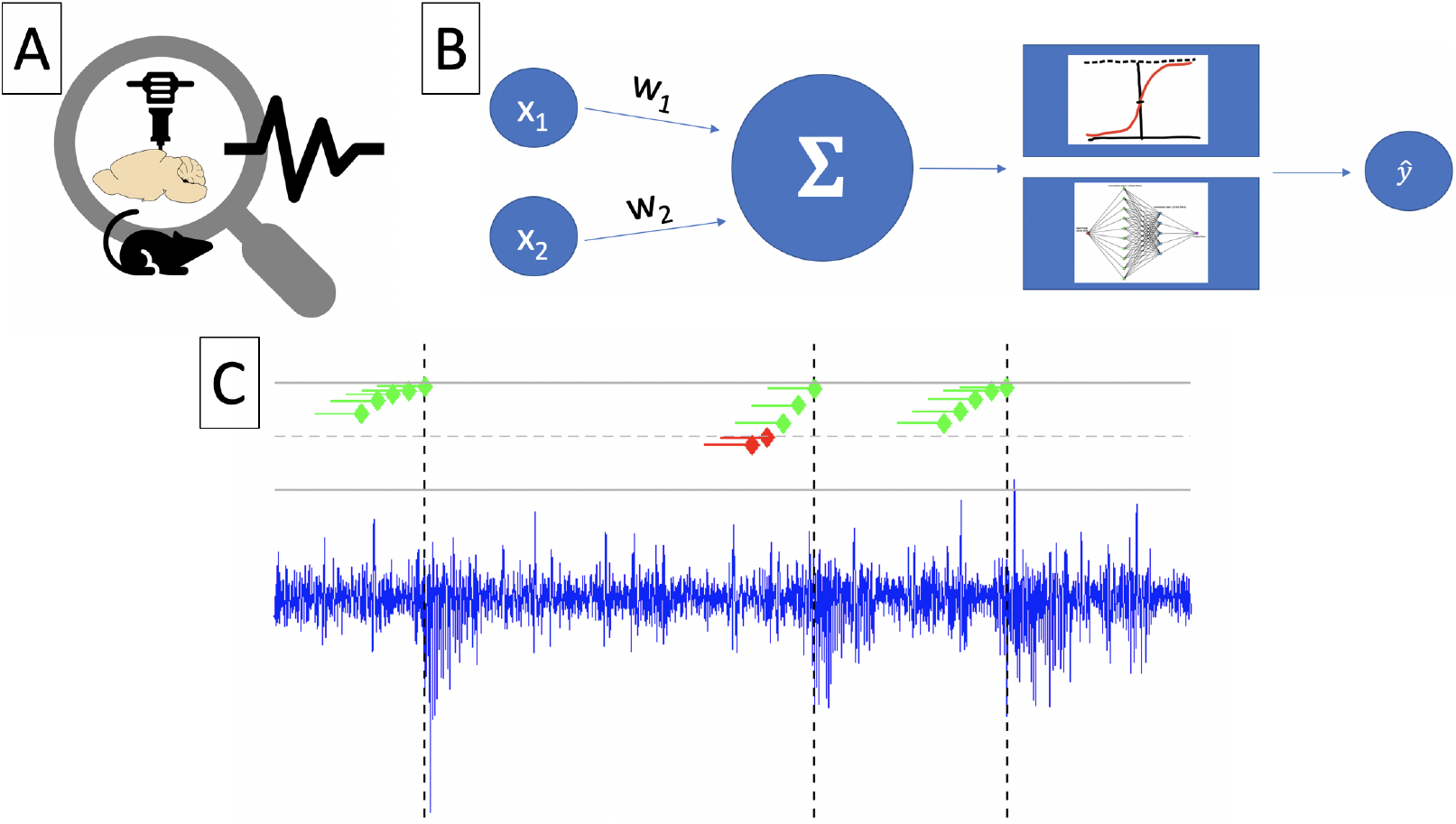
Pipeline detailing approach to seizure prediction using logistic regression. **A.** Cartoon experimental setup detailing Scn8a(+/−) mouse model and recording to signal. **B.** Logistic regression and two-layer convolutional network pipeline with two inputs (1 averaged thalamic and 1 averaged cortical channel from EEG) **C.** Final seizure prediction system showing correct predictions in green, and incorrect predictions in red. Seizure start times marked by black dotted line.

## 2 Related Work

Previous work with general seizure prediction has focused on a binary prediction approach. Petrosian et al.[9] was one of the first studies that focused on investigating the existence of a preictal stage before the seizure through the use of wavelet transformations and found that they were able to detect the presence of preictal stages before the occurrence of the seizure. These findings opened up the way for future predictive methods and studies based on classification with the preictal period. A study done by Li et al.[10] in 2013 showed that seizures could be predicted up to 10 seconds prior to seizure onset with sensitivity of 0.758, despite previous studies basing prediction off of much longer timescales. This allowed future studies to be based off of shorter time scales, which helped simplify the prediction problem and amount of data.

Two additional studies in 2017 used support vector machines to classify EEG seizure signals [11] [12]. One of the studies, Sharif et al. from 2017, was able to achieve sensitivities between 91.8-96.6% [11]. The other study, Direito et al., was the first study to implement a realistic seizure prediction approach by using multi-channel high-dimensional datasets as opposed to the typical dimensionality reduction techniques used in previous papers [12]. By using multiclass support vectors machines on 1206 seizures, Direito et al. was able to achieve sensitivity of 38.47%. While this number was rather low, it showed that seizure prediction could be applied to realistic clinical datasets that have much more data than previous studies. This gave our group the confidence to try and use a large, and realistic EEG dataset with 1913 seizures. Finally, Tsiouris et al. was one of the first to use long short-term memory (LSTM) networks for EEG signal prediction and was able to achieve sensitivity over 99% [13]. We build on these results and attempt to focus on achieving good performance with our models, while also testing how varied the preictal window can be.

This paper specifically focuses on absence seizures. Absence seizures have traditionally been thought of as completely unpredictable events with no defined correlation between different network states [1]. Behaviorally, absence seizures are characterized by periods of quiet wakefulness, when delta waves, brain wave oscillations recorded in an EEG between 1-4 Hz, are most prevalent in the brain [14]. Absence epilepsy is characterized by abnormally synchronous electrical activity within two mutually connected brain regions, the thalamus and cortex [15]. This absence seizure dataset our group has allowed us to explore novel approaches for a difficult type of seizure prediction.

## 3 Methods

### 3.1 Data Collection

Data recording: The data analyzes comes from extra-cellular recordings of individual cortical and thalamic neurons from rodents with genetic absence epilepsy. Local field potential (LFP), a signal analogous to the elecrocorticogram (EEG), was obtained from the Scn8a(+/−) murine model of absence epilepsy via a multi-electrode silicon probe implanted within the somatosensory cortex. Electrodes contacts were 10×10 um in size and gold-plated to achieve an impedance of 250 kOhms. Raw LFP was bandpass filtered between 1-7000 H, sampled at 30 kHz, and subsequently downsampled to 500 Hz for offline analysis. Recordings were performed between the hours of 8 AM and 4 PM in a quiet, isolated room with dim ambient light using an Intan RHD-128 headstage connected to an OpenEphys acquisition board (https://open-ephys.org/acq-board). For more details, see [CITE SOROKIN et al. 2020]. (http://intantech.com/files/Intan_*R*_*HD*2000_1_28*_c_hannel_h_eadstage.pdf*)//

### 3.2 Models

#### 3.2.1 Logistic Regression

Logistic regression is a common statistical modeling tool used for binary classification tasks. The probability of dichotomous outcome event given by *P*_1_ (seizure or non-seizure). In our project, the two classes were defined as epileptic EEG signals and non-epileptic EEG signals (thus referred to as preictal and interictal signals). This probability *P*_1_ is related to a set of explanatory variables that follows the relation shown below.

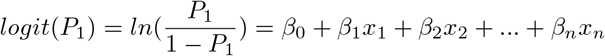

In this equation, we can note that *β*_*n*_ represents the coefficient associated with the explanatory variable *x*_*n*_. Class membership probability is computed via the following relation:

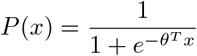

In the above equation, the model will learn the vector parameter, *θ*, and output a value that represents the model confidence in classification grouping. The probability of the dichotomous outcome event is based on a set of explanatory variables and yields a value that is restricted to a binary, two-choice decision such as seizure/non-seizure, mathematically represented as a 1 vs 0 case scenario. Maximum liklihood estimation (MLE) is used to estimate these coefficients *β*_*n*_, and is commonly used in logistic regression. The MLE method is normally chosen since it maximizes the log likelihood which reflects the probability of a successful prediction of the dependent variables. MLE is also an iterative algorithm, which starts with an arbitrary estimate of the coefficients (*β*_*n*_), and minimized using Newton’s method until convergence criterion is reached. We choose to use logistic regression rather than ordinary linear regression (OLR) because logistic regression does not assume an underlying linear relationship between the explanatory variables and response variable. In addition, OLR requires Gaussian distributed independent variables.

#### 3.2.2 Convolutional Neural Networks

Artificial Neural Networks (ANNs) are a common machine learning method that employ artificial neurons or nodes to learn salient trends within the data in order to inform a prediction. ANNs vary in terms of their structure based on the type of problem that they are being implemented for. A relevant category of ANNs are convolutional neural networks (CNNs) that are particularly useful for extract distortion-invariant patterns. Convolutional networks are mostly effective with image recognition, and commonly used to predict seizures by collapsing the raw EEG into a matrix representation. These matrix representations maintain only the relevant information, such as power or frequency extracted from the EEG, and are used in many binary classification seizure prediction methodologies. We use an EEG window length of 3 seconds. In this paper, we adopted a shallow 2-layer CNN model to classify between the two categories of epileptic states. There are 2 blocks that have a convolutional unit. We chose to use a rectified linear activation function (ReLU) that will help overcome the vanishing gradient, as ReLU will not saturate.

### 3.3 Feature Selection

Multichannel epileptic EEG signals are used in this methodology, with 10 chosen cortical and 10 chosen thalamic channels. Feature extraction helps reduce dimenionsality of large datasets, and extract useful information for predictions. Each seizure clip has been segmented into a 12 second window, and the seizure start is aligned at the 8 second mark.

The seizure start was detected by spike sorting software written by author Jordan Sorokin. A preictal length of 3 seconds was chosen for this analysis by checking logistic regression prediction performance averaging different sized preictal windows. These time-dependent logistic regression models were independently trained on their time-specific data. For example, the offset −3 classifier was trained on preictal segments of 3 seconds that ranged from −6 to −3 seconds before seizure onset. The motivation of training time-dependent LR classifiers was to identify valid thresholds for the preictal and interictal cutoffs. Results of these time-dependent classifiers are shown below.

**Figure 2:**
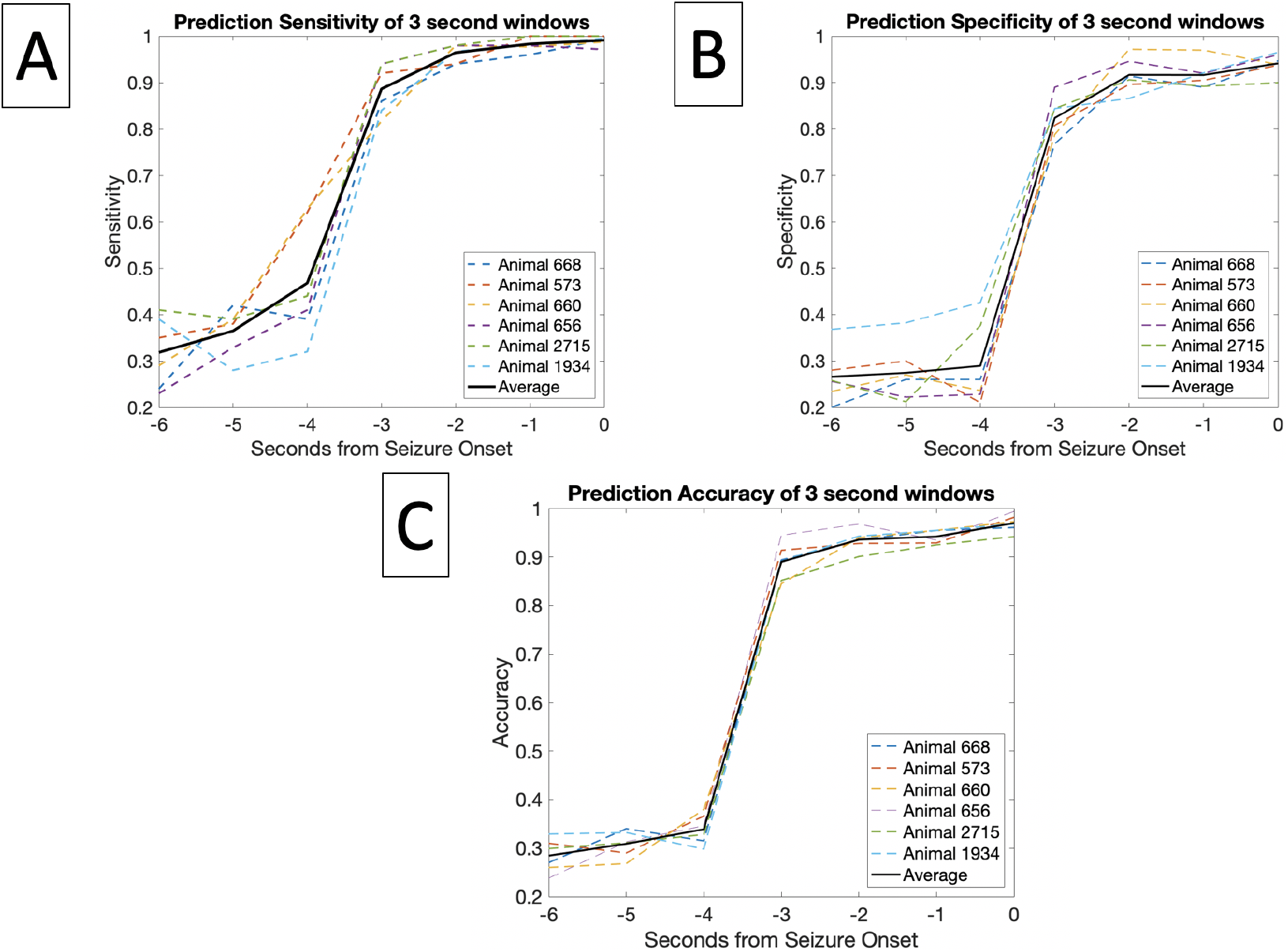
Performance of 6 animals (668, 573, 660, 656, 2715, 1934) and average of time-dependent logistic regression seizure prediction model at 7 different seizure offset times (−6 to 0 seconds before seizure start). Each individual animal trial is shown with a colorful dotted line, and the 6 animal average is shown with a solid black line. **A.** Graph detailing sensitivity 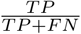. **B.** Graph detailing specificity 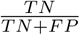. **C.** Graph detailing accuracy 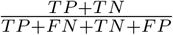

By segmenting seizures into these 12 second blocks, it allows for a more standardized training approach that can be widely applied to any animal with marked seizures. It also allows for easier observation of trends both overall in the dataset, and in particular animals or subsets of recordings.

EEG feature extraction varies between prediction models, and can range from univariate to multi-variate features. Univariate features from EEG include any features regarding one variable, such as information from a single recording site. Multivariate features focus on information from multiple recording sites. For this study, features were extracted from 10 averaged thalamic channels, and 10 averaged cortical channels taken from a 64 channel EEG. Absence epilepsy has been shown to heavily involved the thalamocortical circuit, so only thalamic and cortical channels were chosen for feature extraction [1].

As input to the logistic regression binary classifier and CNN, different features were fed as inputs. For variations of the logistic regression classifier, the inputs consisted of the fast Fourier transform (FFT) of the EEG signal applied to decompose the wave from the time domain into its corresponding frequency signals.

For the convolutional neural network approach, we extracted features of the multichannel EEG by characterizing the energy variation of signals. We de-noise these signals through wavelet transform, and then analyze the power spectrum density (PSD) and use these two-dimensional images as input to our CNN model. These power spectrum density energy diagrams are representations of relative energy intensity of different frequency bands. The PSD analysis is implemented on 3 second frames of the EEG signals, with non-trivial differences noted between the preictal and interictal states.

## 4 Results

### 4.1 Logistic Regression Results

The goal of the logistic regression model approach was a straightforward binary classification using logistic regression, a simplistic model that is not computationally time-intensive. Using a preictal window of 3 seconds, and an interictal window of 5 seconds, the results of the logistic regression binary seizure classification task can be given below. We first show the results of the time-agnostic classifier, which means that the single model was training on aggregated data.In the first table, the 9 animals were kept separate in both training and testing of the model, while in the second, the 9 animals’ 1500 total seizures were shuffled to provide insight into the generalizability of this performance task. A train:test split of 80:20 was chosen for this task on account of the binary classes being balanced and there not being a shortage of data available.

| Logistic Regression Unique Animal Model Results |  |  |  |  |  |  |  |  |
| --- | --- | --- | --- | --- | --- | --- | --- | --- |
| Animal | # of Seizures | TP | FN | TN | FP | Sensitivity(%) | Specificity(%) | Accuracy(%) |
| 668 | 151 | 150 | 1 | 143 | 8 | 99.34 | 94.70 | 97.02 |
| 573 | 176 | 176 | 0 | 182 | 18 | 100 | 93.75 | 96.88 |
| 660 | 246 | 243 | 3 | 231 | 15 | 98.78 | 93.90 | 96.34 |
| 656 | 175 | 170 | 5 | 168 | 7 | 97.14 | 96 | 96.57 |
| 2715 | 330 | 330 | 0 | 297 | 33 | 100 | 90 | 95 |
| 1934 | 202 | 201 | 1 | 195 | 7 | 99.50 | 96.53 | 98.02 |
| 1931 | 87 | 87 | 0 | 87 | 0 | 100 | 100 | 100 |
| 3906 | 77 | 76 | 1 | 77 | 0 | 98.70 | 100 | 99.35 |
| 3908 | 57 | 57 | 0 | 54 | 3 | 100 | 94.74 | 97.37 |
| Logistic Regression Combined Animal Model Results |  |  |  |  |  |  |  |  |
| Animal | # of Seizures | TP | FN | TN | FP | Sensitivity(%) | Specificity(%) | Accuracy(%) |
| Average | 1500 | 1492 | 8 | 1417 | 83 | 99.47 | 94.47 | 96.97 |

We can see that in the table above, each animal-specific model had higher sensitivity than specificity, with the exception of animal 3906. This shows that in each animal-specific case, the model was able to correctly classify more seizures than non-seizure activity. For the combined animal model, this also holds true, with a sensitivity of 99.47% compared to a specificity of 94.47%. This binary prediction task performed well at classifying all forms of seizures and non-seizures overall having a classification accuracy of 96.97% in the generalized model, showing that a binary prediction is feasible with this pipeline.

Additionally, we can see a closer look at the time-agnostic model below with the 0.1s Step Accuracy graph. The graph shows the prediction accuracy of 6 different animals (animals selected based on having > 100 seizures), for 0.1 second steps with a single time-agnostic best-performing LR classifier.

| Convolutional Neural Network Results |  |  |  |  |  |  |  |  |
| --- | --- | --- | --- | --- | --- | --- | --- | --- |
| Seconds offset | # of Seizures | TP | FN | TN | FP | Sensitivity(%) | Specificity(%) | Accuracy(%) |
| 0 | 1500 | 1497 | 3 | 1459 | 41 | 99.8 | 97.27 | 98.53 |
| 1 | 1500 | 1495 | 5 | 1460 | 40 | 99.67 | 97.33 | 98.5 |
| 2 | 1500 | 1478 | 22 | 1444 | 56 | 98.53 | 96.26 | 96.5 |
| 3 | 1500 | 1421 | 79 | 1392 | 108 | 94.73 | 92.8 | 93.77 |
| 4 | 1500 | 1388 | 112 | 1303 | 197 | 92.53 | 86.87 | 89.7 |
| 5 | 1500 | 956 | 544 | 630 | 870 | 63.73 | 42 | 52.87 |
| 6 | 1500 | 562 | 938 | 498 | 1002 | 34.47 | 33.2 | 35.33 |

For the convolutional neural network approach we show a generalized time-agnostic model that has been offset by various increments. We trained the CNN by sliding it along in 1 second increments as shown in the chart, and then 0.1 second increments as shown in the graph. We can see that the convolutional neural network approach reports similarly high accuracies and metrics as the logistic regression model. Additionally, the convolutional neural network approach was an aggregated approach and done on all the animals, giving an idea about how this model would generalize past the animal-specific models like LR. Below the chart we can see a similar 0.1 second step graph showing the performance of the time-agnostic classifier on 0.1 step prediction windows for the CNN. The graph shows the result of the aggregated total 9 animal data, containing 1500 total seizures.

## 5 Discussion

A number of metrics are used to evaluate seizure prediction performance such as the true positive rate (TP) which is defined by the number of EEG segments that are correctly classified as preictal segments, true negative rate (TN) which is defined by the number of EEG segments that are correctly classified as interictal, false positive rate (FP) which is defined as the number of EEG segments that are incorrectly classified as preictal, and the false negative rate (RN) which is defined as the number of segments incorrectly classified as interictal. When evaluating performance of models, we consider these quantitative metrics, as well as overall generalizability of model. Observe the table below for a comparison of our models with other seizure prediction results from the field.

| Study Overview |  |  |  |  |  |
| --- | --- | --- | --- | --- | --- |
| Study | Dataset | Classifier | Sensitivity | Specificity | Accuracy |
| Li[7], 2007 | Absence epilepsy (GAERS) dataset | custom permutation entropy algorithm | 54 | 98.6 | - |
| Maksimenko[4], 2017 | Absence epilepsy (WAG/Rij) dataset | custom SWD algorithm | - | - | 88 |
| Chu[8], 2017 | CHB-MIT dataset | Fourier Transform coefficients and PSD | 77.78 | 0.608 | - |
| Truong[16], 2018 | Freiburg, CHB-MIT, Kaggle | CNN | 75-81.4 | 94-79 | - |
| Rajaguru[17], 2017 | own human EEG dataset | Logistic Regression | 96.04 | 95.71 | 95.88 |
| Daoud[18], 2019 | own dataset | semi-supervised DL | - | 99.7 | 99.6 |
| Mirowski[19], 2009 | Freiburg Seizure Dataset | CNN | 71 | 100 | - |
| Zhang[20], 2019 | own dataset | Logistic Regression and CNN | - | 92.2 | - |
| Tsiouris[13], 2018 | CHB-MIT dataset | LSTM | 100/99.28 | 99.28 | - |
| Tsiouris[21], 2017 | CHB-MIT dataset | SVM | 85.75 | 85.75 | - |
| Subasi[22], 2004 | own dataset | Logistic Regression | 89.2 | 90.3 | - |
| Iasemidis[23], 2003 | own iEEG dataset | custom STLmax algorithm | 82.6 | 83 | - |
| This study, 2020 | Absence seizure dataset (Scn8a) | Logistic Regression | 99.47 | 94.47 | 96.97 |
| This study, 2020 | Absence seizure dataset (Scn8a) | CNN | 99.8 | 97.27 | 98.53 |

Both the Logistic Regression model and the CNN model were able to achieve remarkably good performance. For the Logistic Regression model with short-term fourier transform (FT) features, we can see that the predictive algorithm was able to differentiate between seizures and non-seizures with a 97% accuracy and 99% sensitivity. Contextually, this means that for every 100 events measured, the model was able to correctly identify 97 of them, and out of every 100 seizure events measured, the model was able to correctly identify 99 of them. The CNN model was able to achieve accuracies around 98% with the PSD approach. Compared to traditional seizure prediction literature, we can see that the logistic regression model presented in this paper performed better than most other LR studies, and comparable to more advanced classification models. Subasi et.al.[22] achieved specificity of 90.3% and sensitivity of 89.2% with a similar LR approach, while Mirowski et.al.[19] was able to achieve 100% sensitivity in only 11 out of the 21 patients studied. Compared to other, more complex machine learning classifiers, the LR approach gave comparable results, and the CNN result was able to easily match the current results. A convolutional neural network (CNN) was used by Zhang et.al.[20] to predict seizures using common spatial patterns of EEGs, and achieved a sensitivity of 92.2%. Daoud et.al.[18] employed a semi-supervised approach to raw EEG with a number of different deep learning models such as multilayer perceptron, CNNs and bi-directional LSTMs, as well as fused versions of these models. They achieved high values, namely with their fused bi-LSTM models, around 99.7% for sensitivity and 99.6% for accuracy. In addition, Tsiouris et.al.[13] was also able to achieve remarkably high numbers using an LSTM approach for binary prediction, getting sensitivities around 99-100%. One drawback to using such a powerful model is run-time drastically increases for more computationally intensive models such as LSTMs. This same group employed the less computationally intensive models for an approach with Support Vector Machines (SVMs), and Decision Trees and achieved sensitivity measures at 86.75% for the SVM and 82-83% for the Decision Trees [21]. Our model in comparison utilizing a more basic logistic regression classifier and can achieve sensitivity measures ranging from 97-100%. We can see that the high sensitivity rate indicates that the model is better at predicting seizure-like events, which when translated to a real-life implementation would make for a more effective device.

Additionally, the comparison of the time-dependent LR models with the time-agnostic LR models showed that while specialized training can result in higher prediction accuracies for longer offsets, it is not necessary for a strong prediction. Figure 3 shows that a time-agnostic classifier can achieve high prediction (> 0.9) even when training is highly generalized. While the overall results might give slightly lower metrics than a time-dependent, specialized model, the time-agnostic model is easier to train and implement, and gives comparable results. If this seizure prediction system were to be applied to a closed-loop system with real-time EEG data, then a single time-agnostic classifier would be highly beneficial for ease of training and implementation. Additionally, our inclusion of the convolutional neural network model showed strong performance from the [−3.2 → −2] second interval compared to the time-agnostic best logistic regression model. The CNN model was able to achieve 0.9 accuracy, while the LR model ranged from 0.6 − 0.88 This demonstrates that the CNN model was able to use latent features in this period of time to outperform the LR model, which could indicate that different or fused prediction models would perform the best for clinical implementation.

**Figure 3:**
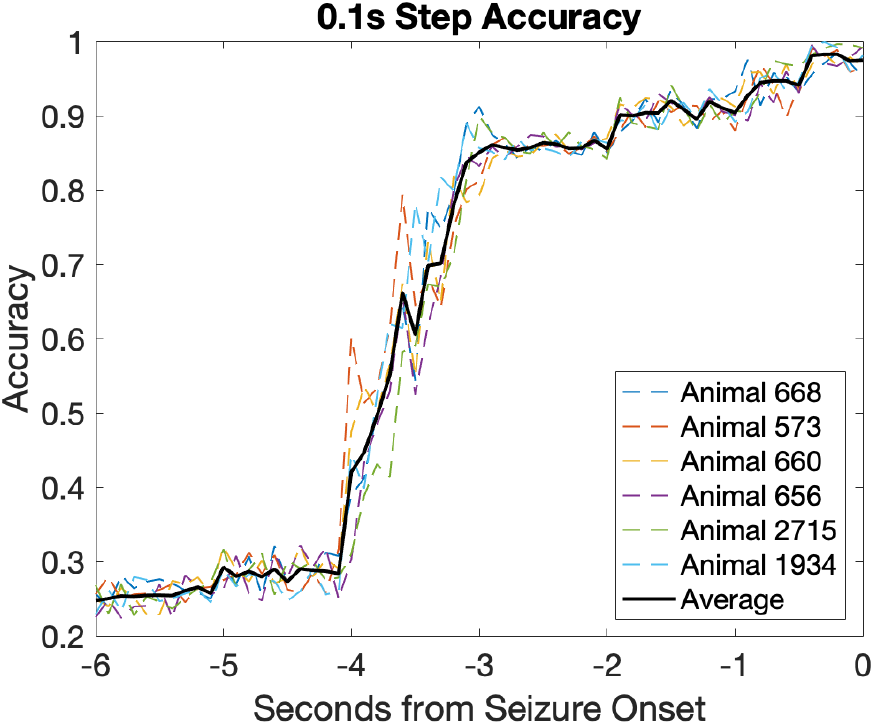
Graph detailing 0.1 second step accuracy for time-agnostic LR models. Prediction accuracies given by the relation 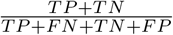 were calculated for each 0.1 step following (6, 5.9, 5.8 … 0.1, 0) seconds before seizure start. Individual performance of 6 animals (668, 573, 660, 656, 2715, 1934) shown in colorful dotted lines, while the 6 animal average is shown with a solid black line.

**Figure 4:**
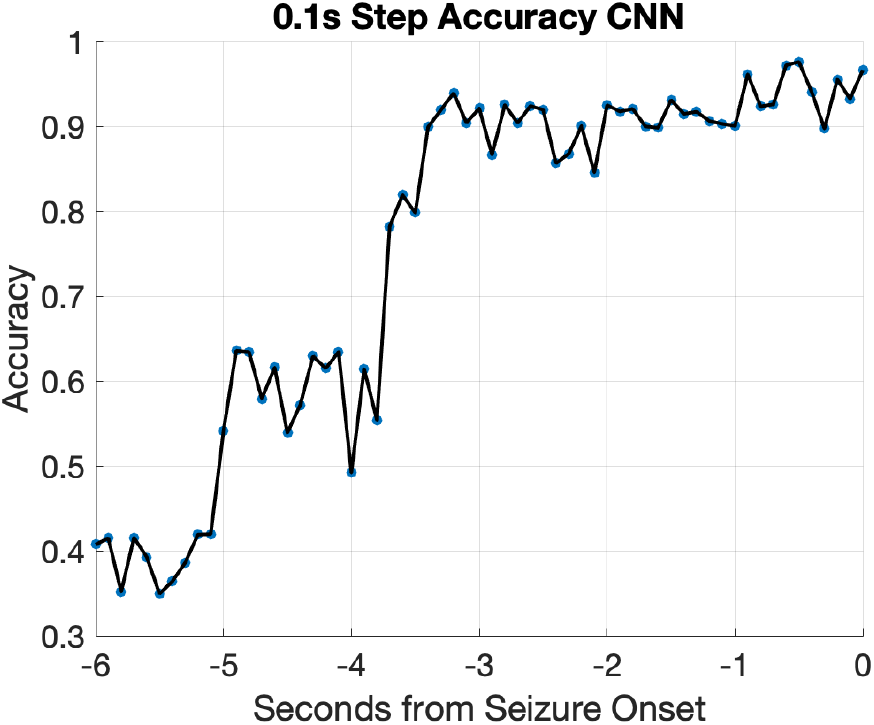
Graph detailing 0.1 second step accuracy for convolutional neural network model. Prediction accuracies given by the relation 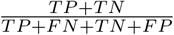 were calculated for each 0.1 step following (6, 5.9, 5.8 … 0.1, 0) seconds before seizure start. Values of the predictions are marked with a blue dot on the graph, while the overall trend is connected with a black line.

## 6 Conclusion

As new machine learning methods are developed, the future of seizure prediction will continue to evolve and change. In this paper we were able to show a two-model approach to an absence seizure prediction pipeline using logistic regression and convolutional neural networks. The logistic regression seizure prediction model achieved accuracies of 97%, and sensitivities of 99%, while the CNN model achieved accuracies of 98% and sensitivities of 99%. We were also able to show that seizure prediction performance was consistent for a certain threshold of a sliding preictal window. The comparison between a time-dependent classifier and a time-agnostic classifier had highly similar evaluation metrics, with a time-dependent being slightly more accurate. However, while the time-dependent was performed slightly better, the time-agnostic still performed well with accuracies 90% for 3-6 seconds before seizure onset. This demonstrated that a time-agnostic model is viable for future implementation in a real-time, closed-loop system. The proposed methodology could be useful for informing preictal window selection for future prediction devices that utilize EEG. Next steps would focus on clinical, closed-loop testing of this methodology, in addition to the inclusion of different feature selections to create a more robust classifier.

